# aenmd: Annotating escape from nonsense-mediated decay for transcripts with protein-truncating variants

**DOI:** 10.1101/2023.03.17.533185

**Authors:** Jonathan Klonowski, Qianqian Liang, Zeynep Coban-Akdemir, Cecilia Lo, Dennis Kostka

## Abstract

DNA changes that cause premature termination codons (PTCs) represent a large fraction of clinically relevant pathogenic genomic variation. Typically, PTCs induce a transcript’s degradation by nonsense-mediated mRNA decay (NMD) and render such changes loss-of-function alleles. However, certain PTC-containing transcripts escape NMD and can exert dominant-negative or gain-of-function (DN/GOF) effects. Therefore, systematic identification of human PTC-causing variants and their susceptibility to NMD contributes to the investigation of the role of DN/GOF alleles in human disease.

Here we present aenmd, a software for annotating PTC-containing transcript-variant pairs for predicted escape from NMD. aenmd is user-friendly and self-contained. It offers functionality not currently available in other methods and is based on established and experimentally validated rules for NMD escape; the software is designed to work at scale, and to integrate seamlessly with existing analysis workflows. We applied aenmd to variants in the gnomAD, Clinvar, and GWAS catalog databases and report the prevalence of human PTC-causing variants in these databases, and the subset of these that could exert DN/GOF effects via NMD escape.

**Availability and implementation:** aenmd is implemented in the R programming language. Code is available on GitHub as an R package (github.com/kostkalab/aenmd.git), and as a containerized command-line interface (github.com/kostkalab/aenmd_cli.git).

## 1. Introduction

Nonsense Mediated mRNA Decay (NMD) is a well-characterized, evolutionarily conserved quality-control mechanism that is essential for embryogenesis and other developmental processes, and it is known to play a role in human disease.[1] NMD guards against compromised transcripts by affecting their degradation vs. translation, including transcripts with variants that introduce premature termination codons (PTCs). PTC-causing variants where a resulting transcript is subject to NMD can exert loss-of-function (LOF) effects in case of haploinsufficiency, where transcripts from both chromosomes are required for normal protein function. For PTC-harboring transcripts that escape NMD, there are additional possibilities of dominant-negative (DN) or gain-of-function (GOF) effects, where the altered protein may interfere with the wild-type version (DN) or where it can possess an altered molecular function or activity domain (GOF). While molecular mechanisms of DN/GOF effects are generally less well understood compared with LOF effects,[2] they do play a significant role in human disease.[1, 3-12].

Given the potential contribution of PTC variants with NMD escape in causing disease, significant new insights into mechanisms of disease pathogenicity can emerge from annotating PTC-containing transcripts with a prediction about their escape from NMD.[3, 7-25] Using an exon-exon junction complex-dependent model of NMD, a notable fraction of PTC-harboring transcripts is predicted to escape NMD [5]. However, this model only describes about half of NMD-escaping human variation accurately, prompting the development of additional approaches for predicting transcript escape from NMD [1, 16, 22, 26-40]. Nevertheless, and despite the relevance of annotating PTC-causing variants with respect to a modified transcript’s susceptibility to NMD, there is a lack of scalable and accessible software addressing that task comprehensively (i.e., for all types of PTC-causing variants). Therefore, we developed aenmd – a software tool for comprehensive annotation of PTC-causing variant-transcript pairs with (predicted) escape from NMD. aenmd makes use of well-established and experimentally validated rules based on PTC location within a transcript’s intron-exon structure,[34, 36] and it integrates well into existing variant analysis pipelines. In the following, we describe aenmd in more detail and report statistics of NMD escape for PTC-causing variants in the Clinvar [41], gnomAD [42], and NHGRI-EBI GWAS catalog [43] resources.

## 2. Materials and Methods

### 2.1 Annotating escape from NMD

aenmd predicts escape from NMD for combinations of transcripts and PTC-generating variants by applying a set of NMD-escape rules, which are based on where the PTC is situated within the mutant transcript. First, the location of the 5’-most (novel) PTC is determined, and then escape from NMD is predicted by the following five rules [34, 36]: Whether

- the PTC located in the last coding exon (last exon rule),
- the PTC located within *d_pen* bp upstream of the penultimate exon boundary (penultimate exon rule; default: *d_pen = 50*)
- the PTC located within *d_css* bp downstream of the coding start site (css rule; default: *d_css = 150*)
- the PTC located within an exon spanning more than 407bp (407 bp rule)
- the transcript is intronless (single exon rule)

See **Figure 1A**. Distances (in bp) are calculated using the PTC nucleotide closest to the coding start site or exon boundary for the css and penultimate exon rules, respectively; variants are assumed to be left-normalized [44] (aenmd provides this functionality). Variants that overlap exon-intron boundaries or splice regions are not currently analyzed by aenmd. Variant-transcript pairs with a PTC conforming to any of the above rules will be annotated to escape NMD, but results for all rules are reported individually by aenmd; this allows users to focus on subsets of rules, if desired. aenmd is implemented in the R programming language [45], making use of the VariantAnnotation [46] and vcfR [47] packages for importing and exporting variants from vcf files, and the Biostrings [48] and GenomicRanges [49] packages for calculating rules. An index containing all PTC-generating SNVs is pre-calculated for a given transcript set and stored in a trie data structure for lookup, using the triebeard package. For non-SNV variants, alternative alleles for overlapping transcripts are explicitly constructed and assessed. This strategy allows us to assess frameshift variants where a PTC is produced downstream of the variant location, and it accounts for both the size and content of sequence insertions, deletions, and insertion-deletions.

**Figure 1:**
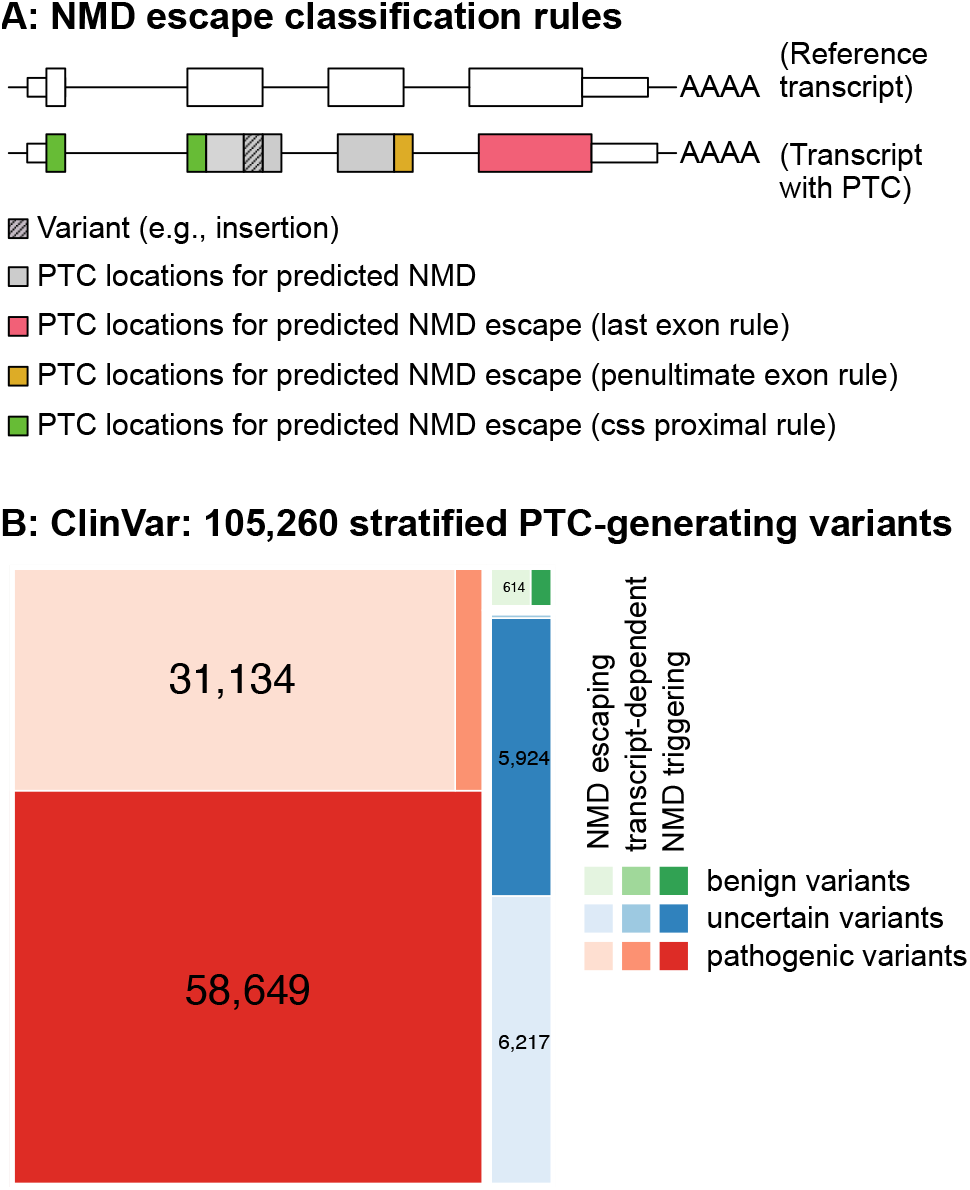
**Panel A** illustrates rules for predicting escape from NMD. Single exon rule, and 407 bp rule are not shown. For the indicated variant, it is impossible for the css proximal rule to apply, because novel PTCs can only be downstream of the variant. **Panel B:** ClinVar variants, stratified by pathogenicity and annotated with predicted escape from NMD. transcript-dependent: the same variant overlaps multiple transcripts and has differing NMD escape predictions.

### 2.2 Data on genetic variants and transcript models

We obtained gnomAD version v2.1.11liftover GRCh38, Clinvar version 20221211, and the NHGRI-EBI GWAS version 20220730, catalog from their respective download sites and annotated variants using aenmd. For our analyses we used transcript models in ENCODE version 105, where we focused on protein-coding transcripts on standard chromosomes that: (a) have an annotated transcript support level of one (or NA for single exon transcripts), and (b) have a coding sequence length divisible by three.

## 3. Results

### 3.1 aenmd R package

The aenmd R-package provides functionality to annotate variant-transcript pairs for predicted escape from NMD within the R ecosystem. Data dependencies (i.e., transcript models) are implemented via specific data packages (see below), and functionality for data import and export (vcf files) is also provided, as is functionality for variant left-normalization. Key differences that set aenmd apart from currently available tools for annotating escape from NMD are: all types of PTC-causing variants (including frameshift variants that do not cause stop codons at the variant site) are annotated, variants are annotated at scale, and differentiated (i.e., rule-specific) output is provided for each transcript-variant pair where NMD-escape rules are applicable. This enables users to focus on the subset of rules most applicable to their situation; for example, some users may choose to focus on “canonical” NMD rules only and choose to ignore the “css proximal”, “single exon”, and “407 bp plus” rules (see **Materials and Methods**). In addition to the R package we also provide a command line interface to aenmd’s functionality.

### 3.2 aenmd_cli command-line interface

We constructed a containerized version of aenmd with all dependencies, which also provides a command-line interface. This allows end-to-end annotation of variants. An input vcf file is read, PTC-generating variants that overlap a specific transcript set (see the aenmd data packages section below) are annotated, and the annotation results are then included in the INFO column of an output VCF file. In this way, the aenmd_cli command line tool makes aenmd easily accessible and its results reproducible; there are no external dependencies, no knowledge of the R programming language is required, and it can be seamlessly integrated into existing variant processing workflows.

### 3.3 aenmd data packages

Annotation for (predicted) escape from NMD is based on the location of a PTC in the context of a transcript model. With aenmd, we provide precompiled annotation packages that provide comprehensive protein-coding transcript sets for the GRCh37 and GRCh38 assemblies of the human genome (data packages: aenmd.data.gencode.v43 and aenmd.data.gencode.v43.grch37, respectively), based on GENCODE version 43 annotations. We also provide a more stringently filtered transcript set based on EN-SEMBL (version 105), containing transcripts with the highest level of transcript support (data package: aenmd.data.ensdb.v105). The aenmd package provides functionality to select between different transcript sets, allowing convenient prediction of NMD escape for GRCh37 and GRCh38 variants.

### 3.4 Annotation of gnomAD, Clinvar and the GWAS catalog

We used aenmd with a high-quality ENSEMBL transcript set (aenmd.data.ensdb.v105 annotation package, see above) to annotate the gnomAD, Clinvar, and GWAS catalog databases of human genetic variation for PTC-generating variants predicted to escape NMD. Our results are summarized in Supplementary Tables S1 - S3. We observe that the fraction of NMD-escape PTC-generating variants varies between 36% (ClinVar), 50% (gnomAD), and 57% (GWAS catalog). The fraction of coding variants in each database that introduce PTCs also varies (10% for ClinVar, 4.1% for gnomAD, and 4.5% for the GWAS catalog). While the absolute number of PTC-generating variants is low for the GWAS catalog (most of its variants are non-coding), we learn from gnomAD that half of the ∼300k PTC-generating variants recovered from ∼125k exome sequences are predicted to escape NMD. Analyzing the ClinVar database (Figure 1B, Supplementary Table S2), we find that for the subset of variants that are considered pathogenic and generate PTCs, 34% (∼31k variants) are predicted to escape NMD. This suggests that escape from NMD may play a substantial role in the disease mechanisms underlying variants of clinical significance annotated in ClinVar.

### 3.5 Comparison with VEP NMD plugin

We note that Ensemble’s Variant Effect Predictor (VEP)[50] provides an NMD annotation plugin that annotates escape from NMD for “stop_gained” variants. This set of variants does not include frameshift variants with a downstream PTC, so the set of variants considered by aenmd and the VEP plugin are inherently different. For example, aenmd annotates ∼200k variant-transcript pairs for ClinVar, while VEP considers ∼77k due to its restrictions on variant type (Supplementary Table S4).

Nevertheless, we systematically compared VEP and aenmd NME escape predictions for the ClinVar database for variants that overlap in transcript set and variant type between the two methods. Overall, we find high consistency of NMD escape predictions (97.5% identical predictions), with 773 (out of 75,840) variant-transcript pairs annotated as NMD escaping by aenmd but not VEP, and with 1,096 pairs annotated as NMD escapeing by VEP but not aenmd. We manually examined a limited set of 20 randomly selected variants with different predictions; results are summarized in supplemental Table S5, and differences are often due to understandable technical differences in the implementation of NMD escape rules.

## 4. Discussion

Here we present aenmd, a self-contained, accessible, and scalable computational tool for annotating (predicted) escape from nonsense-mediated decay (NMD) for variants that generate premature termination codons (PTCs) in a transcript. While we are not aware of a software tool with the same functionality, we compared aenmd to the VEP NMD plugin, which annotates fewer variant types, uses a smaller set of NMD escape rules, and does not report the outcome of individual rules. While NMD predictions were highly consistent between aenmd and VEP, aenmd’s more comprehensive annotations and ease of use allow for better annotation of PTC-generating variants at low computational cost.

## Data and code availability

gnomAD v2.1.1 liftover was downloaded from gnomAD’s downloads website: https://gno-mad.broadinstitute.org/downloads. The Clinvar dataset download, version 2022-12-11 from https://ftp.ncbi.nlm.nih.gov/pub/clinvar/vcf_GRCh38/archive_2.0/2022/. NHGRI-EBI Catalog of human genome-wide association studies, version 2022-07-30, was downloaded from https://www.ebi.ac.uk/gwas/docs/file-downloads. Annotation packages for aenmd are available on GitHub (github.com/kostkalab/aenmd_data.git).

