## supplemental text for "aenmd: Annotating escape from nonsense-mediated decay for transcripts with protein-truncating variants"

Transcript set, prediction rules, and protein-truncating variants

Data generated by aenmd for supplementary tables 1 – 3 utilized aenmd's default settings and transcript set (annotation package aenmd.data.ensdb.v105). Predictions generated by aenmd and VEP's NMD plugin for supplemental tables 4 – 6 utilizes the same NMD escape ruleset as detailed on the Ensembl Blog post for VEP's NMD plugin (https://www.ensembl.info/2022/01/28/cool-stuff-ensembl-vep-can-do-flagging-variants-predicted-to-allow-nmd-escape/): the last coding exon of a transcript, the last 50 bases of the penultimate coding exon of a transcript, the first 100 bases of the coding sequence in the transcript, and intronless transcript. The transcript set for supplemental tables 1 – 6 considered aenmd's transcript set (aenmd.data.ensdb.v105). For all tables, a PTC variant is a variant that causes a PTC in at least one transcript. In tables 1-3 each table is dived into two parts: unique variants and variant-transcript pairs. For supplemental table 4, only variant-transcript pairs are reported.

Transcript-variant pairs vs. unique variants, and canonical and non-canonical rules

In supplemental tables 1 – 4, the values given by unique variants summarizes across variant-transcript pairs. A unique variant is labeled NMD-escaping (or NMD triggering), if NMD escape prediction is consistent across all overlapping transcripts; otherwise, it is labeled transcript-dependent. In tables 1, 3, and 4, transcript-variant pair values consider each pair separately. Canonical variant-transcript pairs are annotated by the exon-exon junction complex dependent NMD escape rules (final exon and penultimate exon rule). Non-canonical refers to annotation by the coding start site proximal, exon >407bp, and the intronless transcript rule. For unique variants, canonical variants are annotated by a canonical rule in any of their overlapping transcripts, non-canonical variants are consistently annotated by non-canonical rules.

Supplemental Table 5

For comparison with VEP, NMD escape rules and parameters used by aenmd were adjusted to match those employed by VEP; variants considered were restricted to those present in both VEP's and aenmd's transcript set.

We considered four separate groups of variant-tx pairs, and examples were sampled randomly, yieldeing 22 variant-transcript pairs:

- Two variant-transcript pairs out of the total 27 variants-transcript pairs (all of which were SNVs) that aenmd called as NMD escaping using the css proximal rule but VEP did not call NMD escaping. (2 variant-transcript pairs)
- One random SNP, two insertions (there are only two) and the only substitution were selected out of 121 total variant-transcript pairs that aenmd called as NMD escaping using the last coding exon rule but VEP did not call NMD escaping. (4 variant-transcript pairs)
- One random SNP, two insertions, two random deletions, and one substitution were selected out of 948 total variant-transcript pairs that aenmd called as NMD escaping using the penultimate exon rule, but VEP did not call NMD. (6 variant-transcript pairs)
- Four random SNPs, three insertions, two random deletions, and one substitution were selected out of 772 total variants that were called NMD escape by VEP but not by aenmd. (10 variant-transcript pairs).
